# An mRNA-LNP-based Lassa virus vaccine induces protective immunity in mice

**DOI:** 10.1101/2023.04.03.535313

**Authors:** Mei Hashizume, Ayako Takashima, Masaharu Iwasaki

## Abstract

The mammarenavirus Lassa virus (LASV) causes the life-threatening hemorrhagic fever disease, Lassa fever. The lack of licensed medical countermeasures against LASV underscores the urgent need for the development of novel LASV vaccines, which has been hampered by the requirement for a biosafety level 4 facility to handle live LASV. Here, we investigated the efficacy of mRNA-lipid nanoparticle (mRNA-LNP)-based vaccines expressing the LASV glycoprotein precursor (LASgpc) or the nucleoprotein (LCMnp) of the prototypic mammarenavirus, lymphocytic choriomeningitis virus (LCMV), in mice using recombinant (r) LCMV expressing a modified LASgpc and wild-type rLCMV. Two doses of LASgpc- or LCMnp-mRNA-LNP administered intravenously or intramuscularly protected mice from a lethal challenge with rLCMVs. Negligible levels of LASgpc-specific antibodies were induced in mRNA-LNP-immunized mice, but robust LASgpc- and LCMnp-specific CD8^+^ T cell responses were detected. Our findings and surrogate mouse models of LASV infection provide a critical foundation for the rapid development of mRNA-LNP-based LASV vaccines.

## Introduction

Rodent-borne mammarenaviruses (*Arenaviridae*: *Mammarenavirus*) include several human pathogens that cause diseases ranging from mild febrile illnesses to life-threatening viral hemorrhagic fever ^1^. Lassa virus (LASV) is a highly prevalent human pathogenic mammarenavirus and the causative agent of Lassa fever (LF), which is endemic in West African countries ^2, 3^. LASV is commonly transmitted to humans from a reservoir in persistently infected rodents, *Mastomys natalensis*, by the inhalation of, direct contact of abraded skin with, or ingestion of, infectious materials contaminated by rodent excreta ^4, 5^. Human-to-human transmission is less common compared with rodent-to-human transmission but can occur in nosocomial settings with poor infection control practices ^6^. LASV is estimated to infect several hundred thousand individuals annually and to be an important public health concern in endemic regions ^2, 3, 7^. During the 2015– 2016 outbreak in Nigeria, the case fatality rates (CFRs) of hospitalized patients with laboratory-confirmed LF were as high as 59.6% ^8^. Despite its significant impact on human health, no licensed vaccines against LASV are available, and current LASV treatment is limited to the off-label use of ribavirin, which may offer partial efficacy with a potential risk of significant side effects ^9, 10, 11^. In addition, environmental changes, including climate change and human population growth in West Africa, may result in an enlarged LASV endemic area ^12, 13^. The lack of medical countermeasures against LASV, exacerbated by recent LASV outbreaks with high CFRs and the expansion of LASV endemic regions, underscores the urgent need to develop novel LASV vaccines.

Similar to other mammarenaviruses, LASV is an enveloped virus with a bi-segmented, single-stranded RNA genome ^1^. The RNA segments, S and L, use an ambisense coding strategy to direct the expression of viral mRNAs from two viral genes arranged in opposite orientations, separated by a noncoding intergenic region (IGR). The S segment RNA encodes the nucleoprotein (NP) and the glycoprotein (GP) precursor (GPC; LASgpc, with particular reference to the LASV GPC). The GPC is co-translationally processed by cellular signal peptidases to generate a stable signal peptide (SSP), and then post-translationally processed by the cellular proprotein convertase subtilisin kezin isozyme-1/site 1 protease (SKI-1/S1P) to generate GP1 and GP2 subunits. GP1 and GP2 form a GP complex with SSP, which is responsible for receptor recognition and cell entry. The L segment RNA encodes a viral RNA-dependent RNA polymerase (L) and matrix RING finger protein (Z). Because cellular immunity has a major role in the recovery and prevention of disease in LF survivor cases and LASV-infected animals, live-attenuated vaccines (LAVs), which can confer long-term cellular and humoral immunity following a single immunization, have been considered a suitable approach for the control of LF ^14, 15, 16^. Consequently, several live-attenuated, virus-vectored LASV vaccine candidates expressing LASV antigens, based on vaccinia virus ^17, 18^, vesicular stomatitis virus (VSV) _19, 20, 21_, ML29 ^22, 23^, and yellow fever 17D ^24, 25^, have been evaluated in LASV animal models, including nonhuman primates (NHPs), with promising results. These studies also reported that GPC and NP were the main protective LASV antigens.

The requirement for highly pathogenic LASV to be handled in a maximum containment (biosafety level 4, BSL-4) laboratory has hampered the development of medical countermeasures against the virus. To study LASV infection under a reduced biocontainment level, a recombinant (r) lymphocytic choriomeningitis virus (LCMV), a worldwide-distributed, prototypic mammarenavirus, whose GPC gene was replaced with wild-type (WT) LASgpc (rLCMV/LASgpc), was generated ^26^. LCMV, thought to be a neglected human pathogen of clinical significance, is a particular threat to immunocompromised or pregnant individuals ^27, 28, 29, 30, 31^. Although rLCMV/LASgpc can be used to investigate LASgpc functions in the context of the natural infection of cultured cells without the need for a BSL-4 facility, the virus is rapidly cleared from C57BL/6 mice _26_. However, K461G, a point mutation in LASgpc that emerged when rLCMV/LASgpc replicated in an HLA-A2.1 transgenic (HHD) mouse ^32^, was observed to prolong viremia by approximately 2 weeks in C57BL/6 mice ^33^. Further characterization of rLCMV/LASgpc mutants revealed that combining V459K and K461G mutations further increased the viral fitness of C57BL/6 mice. Conversely, rLCMV containing corresponding mutations from LCMV GPC toward LASgpc (K465V and G467K) showed poor replication in C57BL/6 mice, indicating that the K465 and G467 of the LCMV GPC are critical for the adaptation of LCMV in mice (*Mus musculus*) ^34^.

mRNA-lipid nanoparticle (LNP)-based coronavirus disease 2019 (COVID-19) vaccines expressing the spike protein of severe acute respiratory syndrome coronavirus 2 (SARS-CoV-2) have been used worldwide and have significantly contributed to the prevention of SARS-CoV-2 infection and progression of COVID-19 to severe disease ^35^. Intriguingly, studies involving humans and animals indicated that COVID-19 mRNA-LNP vaccines elicited neutralizing antibodies as well as spike protein-specific cellular immunity, suggesting that mRNA-LNP-based vaccines are a feasible LASV vaccine modality ^36^. In the present study, we investigated the potential for an mRNA-LNP-based vaccine platform to induce protective immunity in mice, using LASgpc and LCMV NP (LCMnp) as vaccine antigens and rLCMV/LASgpc, containing K465V and G467K mutations (rLCMV/LASgpc^2m^), and WT rLCMV, for challenge experiments.

## Results

### mRNA-LNP vaccines expressing LASgpc (LASgpc-mRNA-LNP) or LCMnp (LCMnp-mRNA-LNP) elicit protective immunity against lethal rLCMV/LASgpc^2m^ infection in C57BL/6 mice

The delivery of *in vitro* transcribed (IVT) mRNA into cells results in the activation of innate immune pathways, which disturb protein production. The use of modified nucleotides, such as pseudouridine and 5-methylcytidine, instead of unmodified nucleotides significantly alleviated the activation of Toll-like receptor (TLR) signaling and protein kinase R (PKR), thereby increasing protein expression in mice ^37, 38^. Importantly, all uridines in the COVID-19 mRNA vaccines produced by BioNTech (BNT162b2) and Moderna (mRNA-1273) are substituted with *N*_1_-methyl pseudouridine ^39^. To evade undesirable innate immune responses and achieve the adequate expression of viral antigens, we substituted all uridines with 5-methoxyuridne, an alternative modified uridine that increased the stability of IVT-mRNA and enhanced protein expression ^40^, to generate 5’-capped and 3’-polyadenylated IVT-mRNAs encoding the LASgpc or LCMnp open reading frame (Supplementary Fig. 1a, b). LASgpc and LCMnp expression in the IVT-mRNA-transfected cells was verified using antibodies specific for LASgpc and LCMnp (Supplementary Fig. 1c, d).

Consistent with previous observations ^33^, we confirmed high levels of viremia (Supplementary Fig. 2a) in C57BL/6 mice 7 days after inoculation with 10^6^ focus forming units (FFU) of rLCMV/LASgpc^2m^ (Fig. 1a). The intracranial inoculation of WT LCMV in mice caused fatal lymphocytic choriomeningitis and the mice died within 8 days of infection ^41^. Next, we investigated whether intracranial (i.c.) inoculation of rLCMV/LASgpc^2m^ to mice also caused a fatal infection. C57BL/6 mice inoculated i.c. with 10^3^ FFU of rLCMV/LASgpc^2m^ all succumbed to the infection albeit with slightly extended survival periods compared with those commonly observed in mice i.c. inoculated with WT LCMV (Supplementary Fig. 2b).

**Fig. 1:**
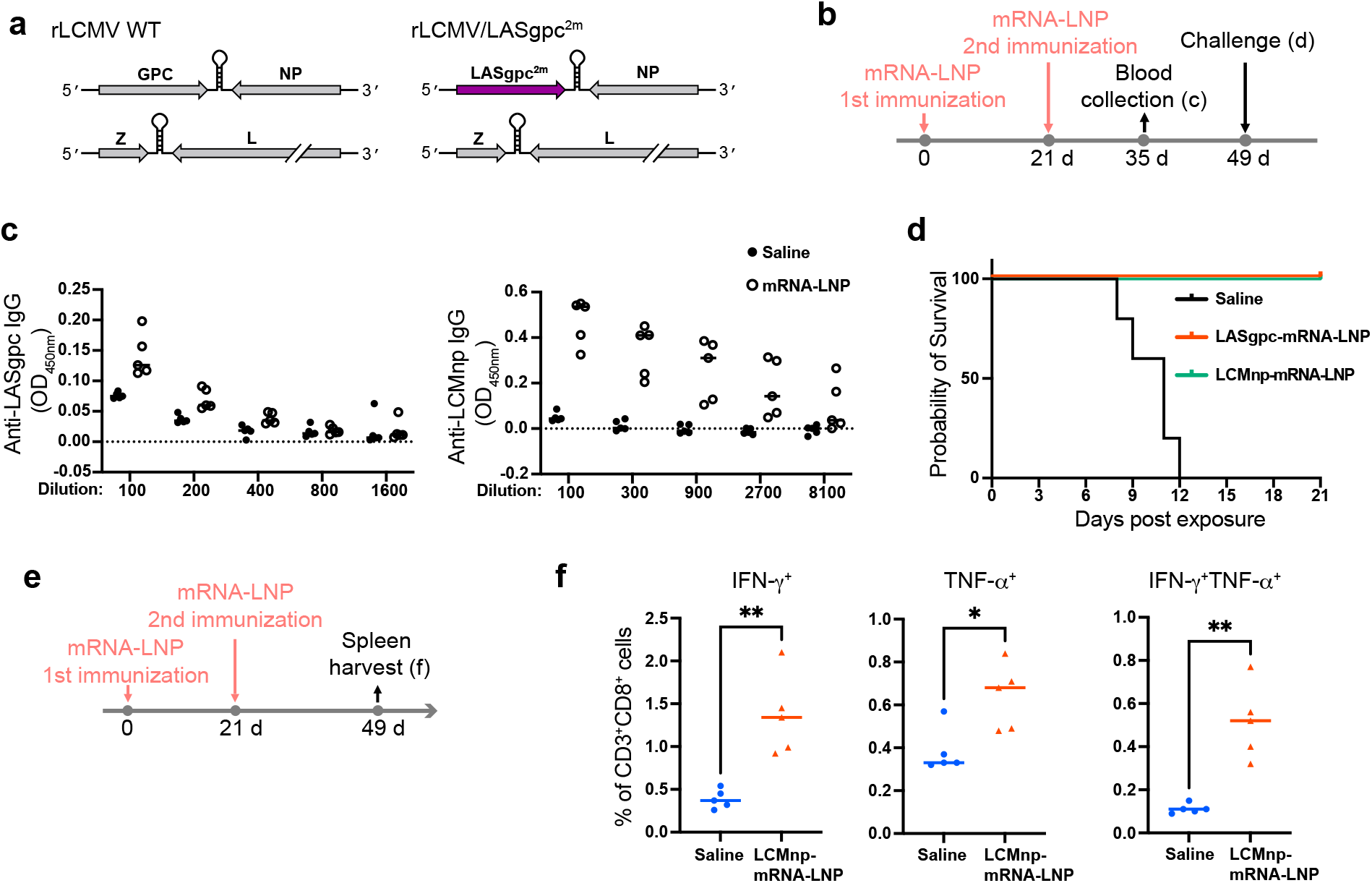
Intravenous immunization of C57BL/6 mice with LASgpc-mRNA-LNP and LCMnp-mRNA-LNP provides protection against rLCMV/LASgpc^2m^. **a** Schematic diagram of the rLCMV/LASgpc^2m^ genome structure. **b–d** Schematic diagram of the experiment (**b**). Six-week-old C57BL/6 mice (n = 5) were immunized i.v. twice with LASgpc-mRNA-LNP or LCMnp-mRNA-LNP, or mock immunized, 21 days apart. At 14 days after the second immunization, anti-LASgpc (left) and anti-LCMnp (right) IgG levels in plasma were determined (**c**). At 28 days after the second immunization, C57BL/6 mice were challenged i.c. with rLCMV/LASgpc^2m^ and survival was monitored daily (**d). e, f** Schematic diagram of the experiment (**e**). Six-week-old C57BL/6 mice (n = 5) were immunized i.v. twice with LCMnp-mRNA-LNP or mock immunized, 21 days apart. At 28 days after the second immunization, splenic CD3^+^CD8^+^ T cells that specifically responded to the NP_396_ peptide were examined for cytokine production by flow cytometry (**f**). The presented data are the mean ± SD. \*\**p* < 0.01, \**p* < 0.05.

To assess the capacity of mRNA-LNP-based vaccines to induce protective immunity, we systemically delivered high doses of mRNA-LNP vaccines intravenously (i.v.) to mice. C57BL/6 mice were immunized twice i.v. with LASgpc-mRNA-LNP or LCMnp-mRNA-LNP incorporating 10 µg of IVT-mRNA, or mock immunized with saline, 21 days apart (Fig. 1b). The levels of LASgpc and LCMnp antibodies in plasma 14 days post-second immunization were examined by ELISAs (Fig. 1c). Consistent with previous reports of virus vector-based LASV vaccine candidates ^42, 43^, we observed negligible levels of LASgpc-specific antibodies (Fig. 1c); however, we confirmed that our ELISA could detect commercially available LASV GP2 antibodies in a dose-dependent manner (Supplementary Fig. 3). Conversely, two doses of LCMnp-mRNA-LNP induced a robust LCMnp-specific antibody response in mice (Fig. 1c). To examine whether two doses of LASgpc-mRNA-LNP or LCMnp-mRNA-LNP conferred protection in mice against lethal virus exposure, mRNA-LNP- or mock-immunized C57BL/6 mice were inoculated i.c. with a lethal dose of rLCMV/LASgpc^2m^ 28 days after the second immunization. As expected, all mock immunized mice succumbed to rLCMV/LASgpc^2m^ infection within 12 days post-virus exposure. By contrast, all mice immunized with LASgpc-mRNA-LNP or LCMnp-mRNA-LNP survived without developing overt clinical signs of disease (Fig. 1d). Low levels of LASgpc-specific antibodies were found in mice immunized with LASgpc-mRNA-LNP, and LCMnp is a cytosolic protein, indicating virus antigen-specific cytotoxic T cell responses might have played a critical role in protection against infection. To assess this, we examined an LCMV-specific CD8^+^ T cell response using a well-defined H-2d-restricted LCMnp T cell epitope peptide (NP_396_). Erythrocyte-free splenocytes obtained from C57BL/6 mice immunized twice with LCMnp-mRNA-LNP- or mock immunized were cultured in the presence of NP_396_ and intracellular cytokine expression levels were examined by flow cytometry (Fig. 1e). Significant increases in IFN-γ- or TNF-α-expressing CD8^+^ T cells from LCMnp-mRNA-LNP-immunized mice were detected, and most TNF-α-positive (TNF-α^+^) cells were also IFN-γ-positive (IFN-γ^+^), indicating the polyfunctional property of antiviral CD8^+^ T cells (Fig. 1f).

### Intramuscular immunization with LASgpc-mRNA-LNP significantly reduces rLCMV/LASgpc^2m^ replication in C57BL/6 mice

Next, we investigated whether protective immunity induced by i.v. immunization of LASgpc-mRNA-LNP could be achieved using a standard intramuscular (i.m.) immunization protocol. C57BL/6 mice were inoculated i.m. twice with LASgpc-mRNA-LNP incorporating 2 µg of IVT-mRNA, or with saline, 21 days apart (Fig. 2a). Consistent with the antibody levels in the plasma from C57BL/6 mice immunized i.v. with LASgpc-mRNA-LNP, LASgpc-specific antibodies were barely detected in the plasma 14 days post-second immunization (Fig. 2b). To evaluate vaccine efficacy by the reduction of virus replication in mice, LASgpc-mRNA-LNP- or mock (saline)-immunized mice were inoculated i.v. with rLCMV/LASgpc^2m^ (10^6^ FFU) 28 days after the second vaccination and virus titers in plasma 7 days post-virus inoculation were determined. I.m. immunization with LASgpc-mRNA-LNP significantly reduced virus titers in the plasma to levels close to or below the lower limit of detection (10^2^ FFU/mL), whereas high virus titers were detected in the plasma from mock immunized mice (Fig. 2c).

**Fig. 2:**
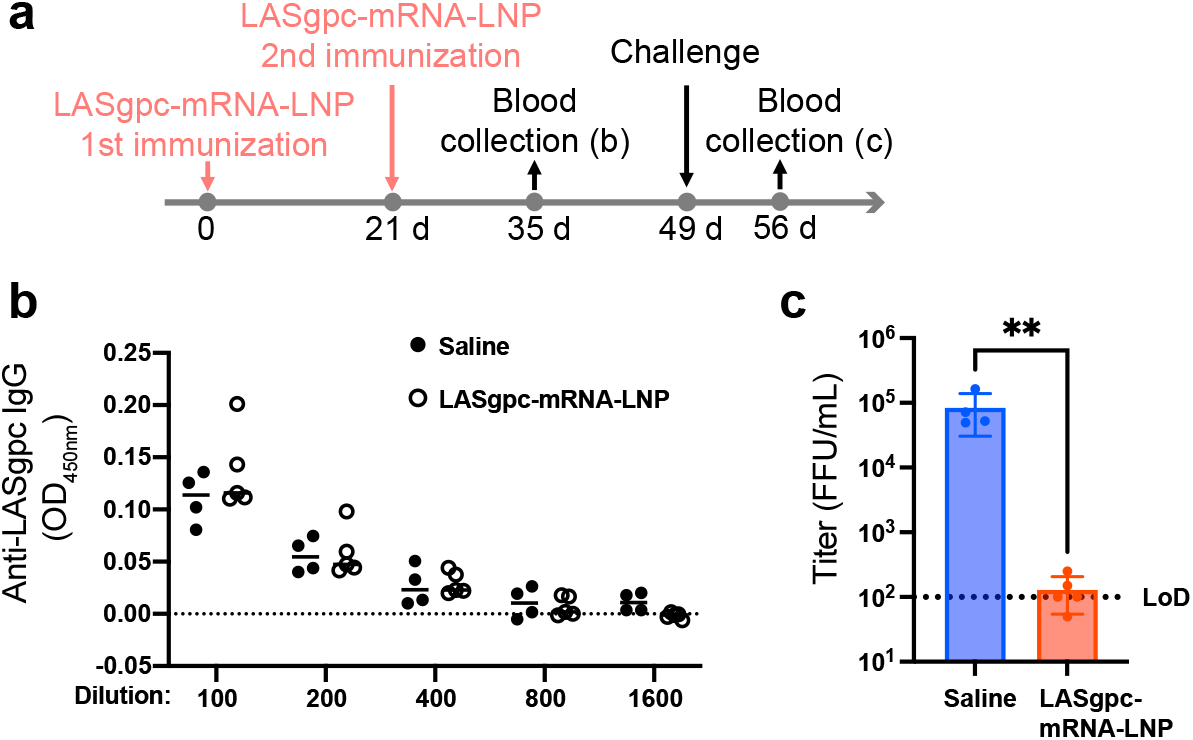
Intramuscular immunization with LASgpc-mRNA-LNP reduces the viral load in C57BL/6 mice. **a** Schematic diagram of the experiment. Six-week-old C57BL/6 mice were immunized i.m. twice with LASgpc-mRNA-LNP (n = 5) or mock immunized (n = 4), 21 days apart. **b** At 14 days after the second immunization, anti-LASgpc IgG levels in plasma were determined. **c** At 28 days after the second immunization, C57BL/6 mice were administered rLCMV/LASgpc^2m^ i.v. and viral titers in plasma were determined. The presented data are the mean ± SD. LoD, low limit of detection. \*\**p* < 0.01.

### Protection against lethal rLCMV/LASgpc^2m^ infection in CBA mice immunized with LASgpc-mRNA-LNP correlates with LASgpc-specific CD8^+^ T cell responses

Immunodominant epitope mapping of LASgpc restricted to H-2k using a 20-mer peptide library identified several peptides that stimulated LASgpc-specific CD8^+^ T cells to produce IFN-γ _44_. To examine whether protection against rLCMV/LASgpc^2m^ infection correlated with the induction of a LASgpc-specific CD8^+^ T cell response, we evaluated the efficacy of LASgpc-mRNA-LNP in CBA (H-2k) mice. CBA mice are highly susceptible to LCMV infection compared with C57BL/6 mice, as evidenced by the finding that a low dose (10^2^ FFU) of WT LCMV (WE strain) inoculated intraperitoneally resulted in 100% mortality in adult CBA mice within 16 days ^45^. To assess vaccine effectiveness, we determined an appropriate challenge dose of rLCMV/LASgpc^2m^. CBA mice were inoculated i.v. with variable doses of rLCMV/LASgpc^2m^ (10^2^–10^5^ FFU per mouse) and clinical signs of disease, body weight changes, and survival were monitored daily. The inoculation of 10^5^ FFU of rLCMV/LASgpc^2m^, but not 10^4^ FFU or less, caused continuous body weight loss until the end of the study (21 days post-virus inoculation) (Fig. 3a), and three of four mice inoculated with 10^5^ FFU of rLCMV/LASgpc^2m^ met the euthanasia criteria (Fig. 3b). Therefore, we selected 10^5^ FFU of rLCMV/LASgpc^2m^ for further studies to assess LASgpc-mRNA-LNP vaccine efficacy in CBA mice.

**Fig. 3:**
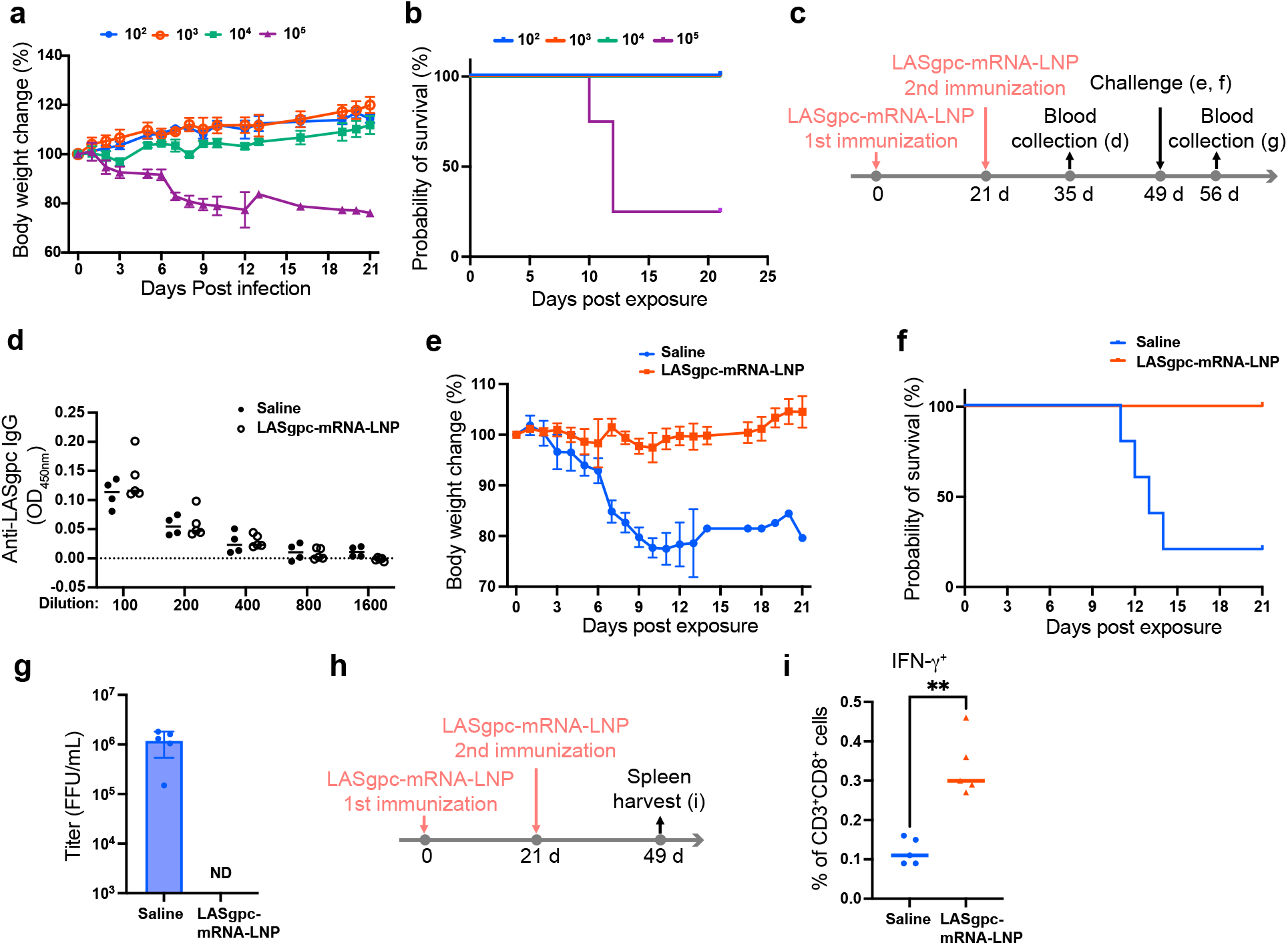
LASgpc-mRNA-LNP immunization provides protection against lethal exposure of CBA mice to rLCMV/LASgpc^2m^. **a**, **b** Eight-week-old CBA mice (n = 4) were inoculated i.v. with rLCMV/LASgpc^2m^. Body weight change (**a**) and survival (**b**) were monitored daily. **c**–**g** Schematic diagram of the experiment (**c**). Six-week-old CBA mice (n = 5) were immunized i.m. twice with LASgpc-mRNA-LNP or mock immunized, 21 days apart. At 14 days after the second immunization, anti-LASgpc IgG levels in plasma were determined (**d**). At 28 days after the second immunization, CBA mice were challenged i.v. with rLCMV/LASgpc^2m^. Body weight change (**e**) and survival (**f**) were monitored daily. Viral titers in plasma were determined at 7 days post-virus exposure (**g**). The presented data are the mean ± SD. ND, not detected. **h**, **i** Schematic diagram of the experiment (**h**). Six-week-old CBA mice (n = 5 per group) were immunized i.m. twice with LASgpc-mRNA-LNP or mock immunized, 21 days apart. At 28 days after the second immunization, splenic CD3^+^CD8^+^ T cells that specifically responded to the LASgpc peptide cocktail were examined for cytokine production by flow cytometry (**i**). \*\**p* < 0.01.

Next, we investigated whether protective immunity against a lethal challenge with rLCMV/LASgpc^2m^ was elicited in CBA mice immunized with LASgpc-mRNA-LNP. CBA mice were immunized i.m. twice with LASgpc-mRNA-LNP incorporating 2 µg of IVT-mRNA, 21 days apart (Fig. 3c). Similar to C57BL/6 mice, barely detectable levels of LASgpc-specific antibodies were observed in the plasma from CBA mice immunized with LASgpc-mRNA-LNP 14 days post-second immunization. At 28 days after the second immunization, CBA mice were inoculated i.v. with 10^5^ FFU of rLCMV/LASgpc^2m^. The mock-immunized CBA mice all had marked body weight loss and four out of the five mice eventually died or met the euthanasia criteria (Fig. 3e, f). By contrast, all LASgpc-mRNA-LNP-immunized CBA mice survived a lethal challenge with rLCMV/LASgpc^2m^ without developing overt clinical signs of disease or body weight loss. Furthermore, LASgpc-mRNA-LNP immunization suppressed the high levels of viremia observed in the mock-immunized mice to below the minimum detection levels (Fig. 3g).

The absence of a strong LASgpc-specific antibody response in CBA mice immunized with LASgpc-mRNA-LNP suggested that a LASgpc-specific CD8^+^ T cell response was involved in the protection. To assess this, we used a LASgpc peptide cocktail to stimulate erythrocyte-free splenocytes collected from CBA mice immunized twice with LASgpc-mRNA-LNP or mock immunized mice, 28 days post-second immunization, and intracellular cytokine expression levels were examined by flow cytometry (Fig. 3h). We confirmed a significant increase in CD8^+^ T cell numbers that produced IFN-γ in response to stimulation with the LASgpc peptide cocktail following LASgpc-mRNA-LNP immunization (Fig. 3i).

### LCMnp-mRNA-LNP confers protection against lethality in a WT rLCMV hemorrhagic fever mouse model

We found that the mRNA-LNP-based vaccine elicited protective immunity against rLCMV/LASgpc^2m^ exposure using two antigens (GPC and NP), two immunization routes (i.v. and i.m.), two virus inoculation routes (i.c. and i.v.), and two mouse strains (C57BL/6 and CBA), strongly suggesting the feasibility of an mRNA-LNP-based vaccine as a potential mammarenavirus vaccine modality. However, although our results demonstrated that an mRNA-LNP-based vaccine can confer protection against severe diseases causing lethality, the infection of C57BL/6 or CBA mice with rLCMV/LASgpc^2m^ may not accurately reflect the pathogenesis of LF. To determine whether an mRNA-LNP-based vaccine could confer protection against an LF-like disease, we used an LCMV mouse model of hemorrhagic disease. I.v. inoculation with 2 × 10^6^ FFU of WT LCMV Clone 13 strain, a derivative of the Armstrong 53b strain, causes acute death (typically within 8 days) associated with thrombocytopenia, coagulation disorder, enhanced vascular permeability, and pulmonary edema in several mouse strains, including FVB, NZO, and NZB ^46, 47, 48^. To assess the efficacy of an mRNA-LNP-based vaccine in a hemorrhagic disease model, FVB mice were immunized i.m. twice with LCMnp-mRNA-LNP incorporating 2 µg of IVT-mRNA, 21 days apart (Fig. 4a). Similar to C57BL/6 mice, FVB mice immunized with LCMnp-mRNA-LNP produced high levels of LCMnp-specific antibodies (Fig. 4b). At 28 days after the second immunization, FVB mice were inoculated i.v. with a typically lethal dose of WT rLCMV and monitored daily for survival and clinical symptoms. Consistent with previous reports ^47, 48^, all mock-immunized FVB mice succumbed to the infection by day 7 post-virus exposure (Fig. 4c). By contrast, FVB mice immunized with LCMnp-mRNA-LNP were fully protected and did not develop any overt clinical signs of disease. To confirm the efficacy of LCMnp-mRNA-LNP in preventing death caused by severe hemorrhagic-like disease correlated with the induction of a robust cytotoxic T cell response, we prepared erythrocyte-free splenocytes from FVB mice immunized with LCMnp-mRNA-LNP (Fig. 4d) and treated them with an H-2q-restricted LCMnp-specific T cell epitope peptide (NP_118_). LCMnp-mRNA-LNP immunization significantly increased the number of CD8^+^ T cells producing IFN-γ, TNF-α, and both IFN-γ and TNF-α, in response to NP_118_ stimulation (Fig. 4e).

**Fig. 4:**
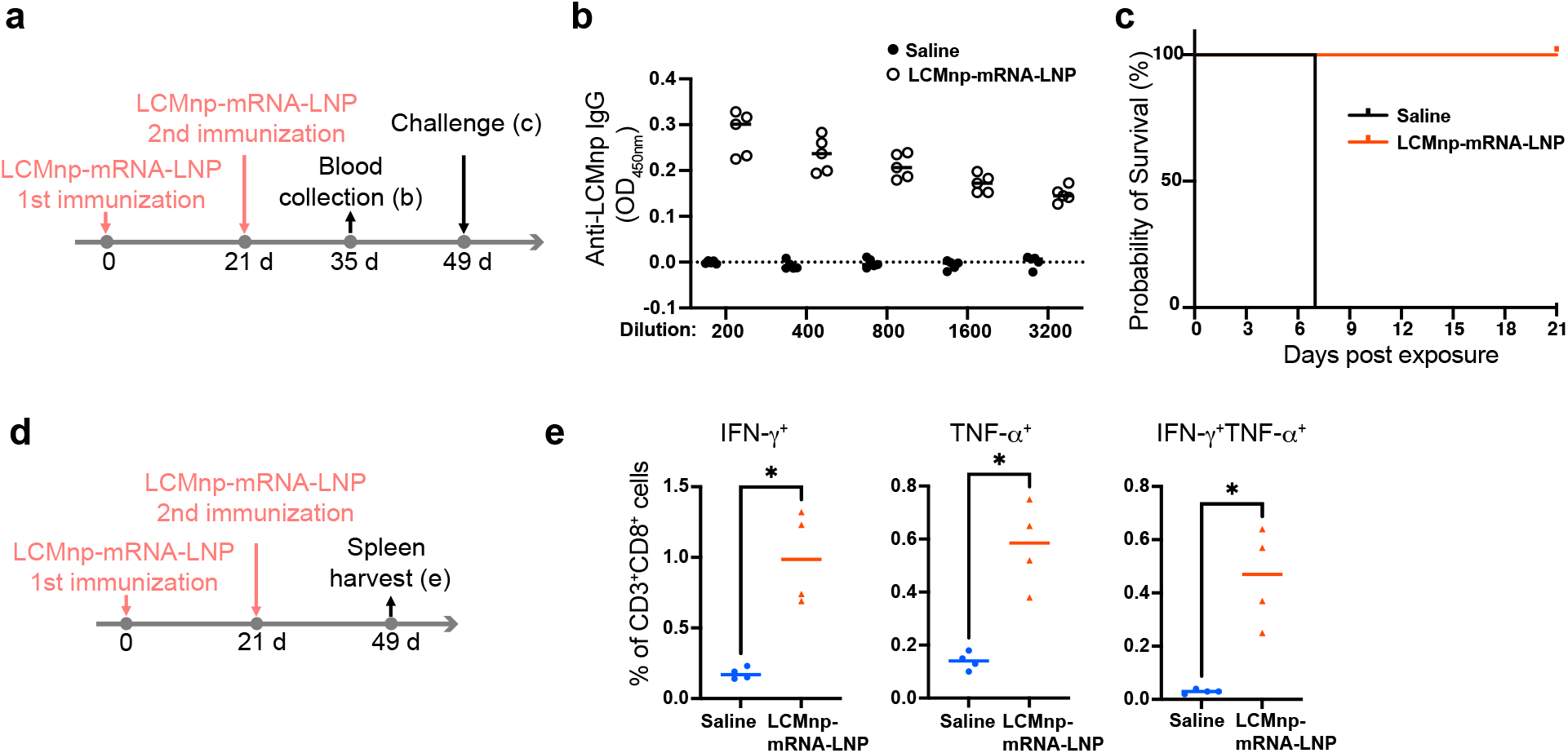
LCMnp-mRNA-LNP immunization provides protection in a hemorrhagic model of FVB mice. **a**–**c** Schematic diagram of the experiment (**a**). Six-week-old FVB mice (n = 5 per group) were immunized i.m. twice with LCMnp-mRNA-LNP or mock immunized, 21 days apart. At 14 days after the second immunization, anti-LCMnp IgG levels in plasma were determined (**b**). At 28 days after the second immunization, FVB mice were challenged i.v. with WT rLCMV and survival was monitored daily (**c**). **d**, **e** Schematic diagram of the experiment (**d**). Six-week-old FVB mice (n = 4 per group) were immunized i.m. twice with LCMnp-mRNA-LNP or mock immunized, 21 days apart. At 28 days after the second immunization, splenic CD3^+^CD8^+^ T cells that specifically responded to the NP_118_ peptide were examined for cytokine production by flow cytometry (**e**). \**p* < 0.05.

## Discussion

LAVs have been the primary modality of LASV vaccine development based on the concept that LASV-specific cell-mediated immune responses have a critical role in the control of LASV, as demonstrated by epidemiological analyses and NHP studies ^14, 43, 49^. In addition, LAVs are effective at establishing herd immunity, which is especially important in developing countries, where achieving optimal levels of vaccination coverage is challenging. However, since the full approval of two COVID-19 vaccines (BNT162b2 and mRNA-1273) by the US Food and Drug Administration, mRNA vaccines are now widely considered an alternative, clinically applicable vaccine modality for infectious diseases. Accordingly, several mRNA vaccines against other viruses, including Zika virus, cytomegalovirus, respiratory syncytial virus, and human immunodeficiency virus, are in phase 2 or 3 clinical trials ^50^. In the present study, we demonstrated the efficacy of mRNA-LNP-based vaccines expressing LASgpc and LCMnp in several surrogate mouse models for LASV infection that correlated with the induction of virus antigen-specific CD8^+^ T cell responses, suggesting that mRNA vaccines constitute an additional vaccine modality against mammarenavirus infections.

LAVs, especially ML29 and an rVSV expressing LASgpc (rVSVΔG-LASV-GPC) ^51^, meet the criteria for optimal vaccine candidates proposed by the World Health Organization (https://www.who.int/publications/m/item/who-target-product-profile-for-lassa-virus-vaccine). The efficacy of a single, low-dose immunization with ML29 has been demonstrated in all available LF animal models, including NHPs, with excellent safety profiles, as highlighted by the finding that ML29 did not cause any mammarenavirus-related diseases in SIV-infected macaques ^52^. The safety of an LAV is particularly relevant in some LASV endemic areas of West Africa that have a high HIV prevalence ^53^. A single immunization of replication-competent rVSVΔG-LASV-GPC conferred complete protection in NHPs against lethal LASV infection ^21^. To be effective, rVSVΔG-LASV-GPC needs to be administered at a high dose (>10^7^ plaque-forming units), which may raise concerns about VSV vector-associated side effects ^54, 55^. LASV causes severe LF in pregnant women, often with fatal outcomes, especially during the third trimester of pregnancy. Moreover, LF causes fetal demise, and fetal evacuation was reported to improve the maternal prognosis ^3^. However, LAVs are not generally recommended for pregnant women (https://www.cdc.gov/vaccines/pregnancy/hcp-toolkit/guidelines.html). COVID-19 mRNA vaccines have been used in pregnant women ^56^ and non-live mRNA-LNP-based LASV vaccines are advantageous as an alternative vaccine candidate for those of limited eligibility to receive LAVs.

A neutralizing antibody (nAb) cocktail against LASgpc exhibited strong therapeutic potential, fully preventing cynomolgus macaques from death, even when administered during the late stage of an LF-like disease ^57^. However, during natural LASV infection, a low titer of nAbs appears late post-infection, indicating the limited contribution of nAbs to protection against acute LASV infection ^58^. Similarly, immunization with ML29 or a vaccinia vector expressing LASgpc resulted in low LASgpc-specific antibody production _43, 44_. By contrast, a single dose of rVSVΔG-LASV-GPC and two doses of a quadrivalent recombinant VSV-vectored LAV candidate, containing one expressing LASgpc, elicited a relatively high LASgpc-specific antibody response, which may have contributed to the protection observed ^19, 21^. In some cases, virion surface protein-binding antibodies induced by viral infection or vaccine inoculation increase virion infectivity and lead to more severe disease ^59, 60^. This phenomenon, called antibody-dependent enhancement (ADE), is one of the concerns associated with the development of vaccines using a virion surface protein as an antigen. In this study, we detected negligible levels of LASgpc-specific antibodies following immunization with two doses of LASgpc-mRNA-LNP, suggesting that the risk of ADE when using LASgpc-mRNA-LNP might be limited. Furthermore, LCMnp-mRNA-LNP induced a robust LCMnp-specific antibody response. Although LCMnp antibodies may not have a significant impact on protection, these antibody levels may be useful to monitor the immune response to the vaccination.

In the present study, we demonstrated that LASgpc-and LCMnp-mRNA-LNP elicited protective immunity in mice against a lethal challenge with rLCMVs. On the basis of these findings, future studies to investigate the effectiveness of mRNA-LNP-based vaccines expressing LASgpc and LASV NP in an LASV infection system are warranted. Findings obtained from studies using LCMV often translate to LASV ^61, 62, 63, 64^. In addition, limited access to BSL-4 facilities together with the high cost of using NHPs are major factors slowing LF vaccine development. Therefore, surrogate small animal models that can be studied in a reduced biocontainment setting are crucial for accelerating the development of LF vaccines, especially at the early stage of research ^65^. Because of the vast resources presently available for physiological, biochemical, immunological, and genetic analyses, a mouse infection model would be particularly useful. CBA (CBA/J) mice that received CD8^+^ T cell-depleted splenocytes from ML29-immunized donors all died after a lethal challenge by the i.c inoculation of ML29, similar to LF infection where T cell-mediated immunity was critical for the control of viral replication ^44^. This CBA-ML29 mouse model also provided information about CD8^+^ T cell epitopes. We found a correlation between the efficacy of mRNA-LNP-based vaccines and the induction of LASgpc-specific CD8^+^ T cell responses. Furthermore, we report that the i.v. inoculation of rLCMV/LASgpc^2m^ caused fatal disease in CBA mice, with gradual body weight loss and a high level of viremia, and that viral replication was controlled by CD8^+^ T cell immunity. This CBA-rLCMV/LASgpc^2m^ mouse model will also allow us to evaluate vaccine efficacy in the context of viral replication, progression of disease, lethality, and cell-mediated immunity. This model will therefore be useful, particularly when investigating various combinations and ratios of LNP components, which may have a significant impact on mRNA delivery efficiency and target cells as well as the immunogenicity of an mRNA-LNP vaccine ^66^, and will contribute to minimizing the workload associated with validation assays in BSL-4 laboratories, facilitating the rapid development of LF vaccines.

## Methods

### Mice

All animal experiments were approved by the Animal Care and Use Committee of the Research Institute for Microbial Diseases, Osaka University. Specific-pathogen-free C57BL/6N and CBA/N mice were purchased from Japan SLC (Hamamatsu, Shizuoka, Japan). Specific-pathogen-free FVB/N mice were purchased from CLEA Japan (Tokyo, Japan). Mice were euthanized humanely at a terminal stage when there was >25% body weight loss or at the study endpoint.

### Cells

HEK293T (American Type Culture Collection, ATCC, Manassas, VA, USA, CRL-3216), HEK293 (ATCC, CRL-1573), and Vero E6 (ATCC, CRL-1586) cells were cultured in Dulbecco’s modified Eagle’s medium (DMEM, Nacalai Tesque, Kyoto, Japan) containing 10% heat-inactivated fetal bovine serum (FBS, Thermo Fisher Scientific, Waltham, MA, USA), 100 U/mL penicillin, and 100 µg/mL streptomycin (Nacalai Tesque) (10% FBS/DMEM) at 37°C and 5% CO_2_. BHK-21 (ATCC, CRL-3216) cells were cultured in 10% FBS/DMEM supplemented with 5% tryptose phosphate broth (Thermo Fisher Scientific) at 37°C and 5% CO_2_.

### Viruses

The two different rLCMVs used in this study were generated by reverse genetics as described previously ^34, 67, 68^. For the generation of WT rLCMV clone 13 strain (WT rLCMV), BHK-21 cells seeded at 7 × 10^5^ cells per well (six-well plate) and cultured overnight were transfected with mPol1Sag Cl-13 (0.8 µg) and mPol1Lag Cl-13 (1.4 µg) that direct the RNA polymerase I (Pol-I)-mediated intracellular synthesis of S and L antigenome RNA species from the LCMV clone 13 strain ^67, 69^, together with pC-NP (0.8 μg) and pC-L (1 μg) that supply the trans-acting factors LCMV NP and L ^70^ using 10 μL of Lipofectamine 2000 (Thermo Fisher Scientific) and incubated at 37°C and 5% CO_2_. At 5 h post-transfection, the transfection mixture was removed and fresh medium was added to the well. After 3 days of incubation at 37°C and 5% CO_2_, the cell culture medium (tissue culture supernatant, TCS) was removed, fresh medium was added to the well, and the plates were cultured at 37°C and 5% CO_2_ for another 3 days. The TCS collected 6 days post-transfection was used to amplify the rescued virus using BHK-21 cells as a cell substrate. The rescue of rLCMV/LASgpc^2m^ was as described for WT rLCMV using pPol1S Cl-13(LASV-GPC/KGGS) ^34^, where the GPC gene of mPol1Sag Cl-13 was replaced with the modified LASgpc gene containing nucleotide substitutions corresponding to the V459K and K461G mutations, instead of mPol1Sag Cl-13.

### mRNA synthesis and formulation

DNA fragments composed of the T7 promoter sequence with a GGG sequence at the 3′ end, the human β-globin (NCBI Accession ID, NM_00518.5) 5ʹ-UTR sequence, either the LASgpc (NCBI Accession ID, NC_004296.1) or LCMnp (NCBI Accession ID, KY514256.1) ORF with a stop codon, and the human β-globin 3′-UTR sequence, in this order, were amplified by PCR. These DNA fragments were used as templates to generate 5ʹ-capped and 3ʹ-polyadenylated mRNAs using the T7 mScript Standard mRNA Production System (Cellscript, Madison, WI, USA), where uridine-5ʹ-triphosphate was replaced by 5-methoxyuridine-5′-triphosphate (TriLink BioTechnologies, San Diego, CA, USA). IVT-mRNAs were then purified using a Monarch RNA clean up kit (New England Biolabs, Ipswich, MA, USA), and the quality of purified IVT-mRNAs was assessed by non-denaturing agarose gel electrophoresis using RNA High for Easy Electrophoresis (DynaMarker Laboratory, Tokyo, Japan).

LNP formulations (IVT-mRNA-LNPs) were prepared from GenVoy-ILM and IVT-mRNA using a microfluidic mixer (NanoAssemblr Ignite, Precision Nanosystems, Vancouver, BC, Canada). The N/P ratio (the molar ratio between cationic amines on ionizable lipids to negatively charged phosphates on mRNA) was set to 4. The resulting samples were diluted with 1× Formulation Buffer 2 (Precision Nanosystems), concentrated by Amicon Ultra centrifugal filters (Millipore, Burlington, MA, USA), and passed through a 0.2-µm filter according to the manufacturer’s recommendations (NanoAssemblr Ignite Training Kit with GenVoy-ILM, Precision Nanosystems). Sterile-filtered mRNA-LNPs were stored at 4°C before use. The concentration of IVT-mRNA incorporated into LNPs was determined using a Quant-iT Ribogreen Assay Kit (Thermo Fisher Scientific).

### Virus titration

rLCMV titers were determined by an immunofocus forming assay as described previously, with minor modifications ^71^. Vero E6 cells seeded in 96-well plates at 2 × 10^4^ cells per well and cultured overnight were inoculated with 10-fold serial dilutions of rLCMV. After 20 h of incubation at 37°C and 5% CO_2_, the cells were fixed with 4% paraformaldehyde in phosphate-buffered saline (PBS) (4% PFA/PBS) (Nacalai Tesque) or neutral-buffered 10% formalin (Fujifilm Wako Pure Chemical Corporation, Osaka, Japan, Wako). Intracellular LCMnp was visualized by an immunofluorescence assay. After cell permeabilization and blocking with 1% normal goat serum (Nacalai Tesque) in dilution buffer [0.3% Triton X-100 (Sigma-Aldrich, St. Louis, MO, USA) in PBS containing 3% bovine serum albumin (Nacalai Tesque)], the cells were incubated with a primary antibody against LCMV NP (VL-4, Bio X Cell, Lebanon, NH, USA), followed by a secondary anti-rat IgG antibody conjugated with Alexa Fluor 568 (anti-rat-AF568) (Thermo Fisher Scientific). Fluorescent images were captured with the CQ1 Confocal Quantitative Image Cytometer (Yokogawa Electric Corporation, Tokyo, Japan), and the NP-positive LCMV-focus numbers were determined using the high-content analysis software, CellPathfinder (Yokogawa Electric Corporation). Virus titers were calculated by multiplying the NP-positive LCMV-focus number by the corresponding dilution factor.

### Assessment of LCMnp expression in cells transfected with IVT-mRNA encoding LCMnp ORF (LCMnp-mRNA)

HEK293 cells seeded in 96-well plates at 5 × 10^4^ cells per well and cultured overnight were transfected with 200 ng of LCMnp-mRNA or mock transfected using 0.5 µL of Lipofectamine 2000 and incubated at 37°C and 5% CO_2_. At 5 h post-transfection, the transfection mixture was removed, fresh medium was added to the wells, and the cells were incubated at 37°C and 5% CO_2_. At 24 h post-transfection, cells were fixed with 4% PFA/PBS and intracellular LCMnp was fluorescently visualized by an immunofluorescence assay, as described in the Virus titration section, using a secondary anti-rat IgG antibody conjugated with Alexa Fluor 488 (Thermo Fisher Scientific) instead of anti-rat-AF568. Fluorescent images were captured with the CQ1 Confocal Quantitative Image Cytometer (Yokogawa Electric Corporation).

### Assessment of LASgpc expression in cells transfected with IVT-mRNA encoding LASgpc ORF (LASgpc-mRNA)

HEK293 cells seeded in 12-well plates at 4.5 × 10^5^ cells per well and cultured overnight were transfected with 1.8 µg of LCMnp-mRNA or mock transfected using 4.5 µL of Lipofectamine 2000 and incubated at 37°C and 5% CO_2_. At 5 h post-transfection, the transfection mixture was removed, fresh medium was added to the wells, and the cells were incubated at 37°C and 5% CO_2_. At 24 h post-transfection, total cell lysates were prepared by lysing the cells with PD buffer (0.5% Triton X-100, 250 mM NaCl, 50 mM Tris-HCl, 10% glycerol, 1 mM MgCl_2_, 1 µM CaCl_2_) containing Halt Protease and Phosphatase Inhibitor Cocktail (Thermo Fisher Scientific). The resulting cell lysate was clarified by centrifugation at 20,600 ×*g* and 4°C for 5 min. The clarified cell lysate was mixed at a 3:1 ratio with 4× Laemmli sample buffer [277.8 mM Tris-HCl, 44.4% glycerol, 4.4% sodium dodecyl sulfate (SDS), 0.02% bromophenol blue] containing 2-mercaptoethanol and denatured at 98°C for 5 min. Protein samples were fractionated by SDS-polyacrylamide gel electrophoresis using a 4%–20% Mini-PRO TEAN TGX Gel (BioRad, Hercules CA, USA), and the resolved proteins were transferred by electroblotting onto polyvinylidene difluoride membranes (Immobilon-P PVDF Transfer Membranes). To detect LASV GP2, the membranes were incubated with a rabbit polyclonal antibody to LASV GP2 (PA5-117438; Thermo Fisher Scientific) and then with horseradish peroxidase (HRP)-conjugated anti-rabbit IgG antibody (Jackson ImmunoResearch Laboratories, West Grove, PA, USA). The Chemi-Lumi One L chemiluminescent substrate (Nacalai Tesque) was used to generate chemiluminescent signals that were visualized with a chemiluminescent imager (Amersham ImageQuant 800, Cytiva, Tokyo, Japan).

### In vivo characterization of rLCMV/LASgpc^2m^

Eight-week-old C57BL/6N or CBA/N mice were inoculated i.v. with rLCMV/LASgpc^2m^ at 10^6^ FFU, unless otherwise indicated, or i.c. with 10^3^ FFU of the virus and monitored daily for signs of disease, body weight, and survival. At 7 days post-inoculation (dpi), blood samples were collected from i.v. virus inoculated mice to determine the viral load in the plasma.

### Immunization with IVT-mRNA-LNP and rLCMV/LASgpc^2m^ challenge of C57BL/6N mice

Six-week-old C57BL/6N mice were immunized i.v. or i.m. twice with IVT-mRNA-LNP, containing 10 µg (for i.v. immunization) or 2 µg (for i.m. immunization) of IVT-mRNA, or mock immunized with saline, 21 days apart. Then, 14 days after the second immunization, blood samples were collected to assess antigen-specific antibody production. At 28 days after the second immunization, mice were inoculated i.c. with 10^3^ FFU of rLCMV/LASgpc^2m^ or i.v. with 10^6^ FFU of the virus and monitored daily for signs of disease and survival. At 7 dpi, blood samples were collected from i.v. virus-inoculated mice to determine the viral load in the plasma.

### Immunization with LASgpc-mRNA-LNP and rLCMV/LASgpc^2m^ challenge of CBA/N mice

Six-week-old CBA/N mice were immunized i.m. twice with LASgpc-mRNA-LNP, containing 2 µg of LASgpc-mRNA, or mock immunized with saline, 21 days apart. Then, 14 days after the second immunization, blood samples were collected to assess LASgpc-specific antibody production. At 28 days after the second immunization, mice were inoculated i.v. with 10^5^ FFU of the virus, and monitored daily for signs of disease, body weight, and survival. At 7 dpi, blood samples were collected to determine the viral load in the plasma.

### Immunization with LCMnp-mRNA-LNP and WT rLCMV challenge of FVB/N mice

Six-week-old FVB/N mice were immunized i.m. twice with LCMnp-mRNA-LNP, containing 2 µg of LCMnp-mRNA, or mock-immunized with saline, 21 days apart. Then, 14 days after the second immunization, blood samples were collected to assess LCMnp-specific antibody production. At 28 days after the second immunization, mice were inoculated i.v. with 2 × 10^6^ FFU of the virus and monitored daily for signs of disease and survival.

### Detection of LCMnp- and LASgpc-specific antibodies by ELISA

Vero E6 cells seeded in six-well plates at 4 × 10^5^ cells per well and cultured overnight were inoculated (multiplicity of infection = 0.01) with rLCMV/LASgpc^2m^, incubated at 37°C and 5% CO_2_ for 48 h, and used to prepare the LCMnp antigen. HEK293T cells seeded in 12-well plates at 4.5 × 10^5^ cells per well and cultured overnight were transfected with 1 µg of a plasmid expressing LASgpc (pCAGGS-LASV-GPC) ^62^, incubated at 37°C and 5% CO_2_ for 24 h, and used to prepare the LASgpc antigen. Cells were harvested, washed with PBS, and lysed in RIPA buffer (50 mM Tris-HCl [pH 7.6], 150 mM NaCl, 1% NP-40, 0.5% sodium deoxycholate, 0.1% sodium dodecyl sulfate) (Nacalai Tesque) containing Halt Protease and Phosphatase Inhibitor Cocktail (Thermo Fisher Scientific). Cell lysates were then sonicated and clarified by centrifugation at 10,000 ×*g* and 4°C for 15 min. Clarified cell lysates were diluted with ELISA Coating buffer (Abcam, Cambridge, UK), coated onto ELISA plates (BioLegend, San Diego, CA, USA) at a final protein concentration of 100 ng/well and incubated at 4°C overnight. Plates were washed six times with PBS-T (PBS supplemented with 0.2% Tween 20) before 300 µL of blocking buffer (PBS-T containing 3% normal goat serum and 2% skim milk) was added and incubated at 37°C for 2 h. Two- or three-fold serial dilutions of plasma were added to the plates, followed by incubation at 4°C overnight. Plates were washed six times with PBS- T before the addition of anti-mouse IgG-HRP (Jackson ImmunoResearch Laboratories) and then incubated at 37°C for 1 h. The plates were washed again with PBS-T and developed with TMB substrate [ELISA POD Substrate TMB Kit (HYPER), Nacalai Tesque]. The reaction was stopped by the addition of stop solution and the optical density was measured at 450 nm using a muti-mode plate reader (SpectraMax iD5, Molecular Devices, San Jose, CA, USA).

### Specific CD8+ T cell responses to LCMnp and LASgpc immunodominant peptides

Six-week-old C57BL/6N, FVB/N, and CBA/N mice were immunized i.m. twice with IVT-mRNA-LNP containing 2 µg of IVT-mRNA, or mock immunized with saline, 21 days apart. At 28 days after the second immunization, spleens were collected from euthanized mice, and erythrocyte-free splenocytes were prepared. Splenocytes from LCMnp-mRNA-LNP immunized mice were stimulated with 2 µg/mL of H-2d (C57BL/6N)-restricted LCMV NP_396_ peptide (Medical & Biological Laboratories, Aichi, Nagoya, Japan, MBL) or H-2q-restricted LCMV NP_118_ peptide (MBL) and 50 U/mL of recombinant mouse IL-2 (BioLegend) for 1 h, and then 4 h in the presence of Protein Transport Inhibitor Cocktail (Thermo Fisher Scientific). Splenocytes from LASgpc-mRNA-LNP-immunized mice were stimulated with the LASgpc peptide cocktail, containing three 20-mer-long peptides (amino acid positions 57–76, 218–237, and 225–244, 10 µM each) and 50 U/mL of recombinant mouse IL-2 (BioLegend) overnight, and then 4 h in the presence of Protein Transport Inhibitor Cocktail (Thermo Fisher Scientific). Cell Stimulation Cocktail (Thermo Fisher Scientific) was used, instead of peptides, to treat cells for 5 h as the positive control. Cells were stained with cell surface marker antibodies (CD3ε: PerCP/Cyanine5.5, clone 145-2C11; CD8a: FITC, clone 53-6.7; both BioLegend) and Fixable Viability Dye eFluor 780 (Thermo Fisher Scientific). After cell fixation with 4% PFA/PBS and permeabilization with Perm/Wash Buffer (BD Biosciences, San Jose, CA, USA), intracellular cytokine staining was performed with antibodies to IFN-γ (PE, clone XMG1.2; BioLegend) and TNF-α (APC, clone MP6-XT22; BioLegend). Flow cytometric analysis was performed using an SH800H flow cytometer (Sony, Tokyo, Japan) and data were analyzed using Cell Sorter Software (Sony).

### Statistics

GraphPad Prism 9 (GraphPad, San Diego, CA, USA) was used for all the statistical analyses. Statistically significant differences were determined by Mann-Whitney test.

## Data availability

All data are available in the main text and the Supplementary Information. Source data are provided with this paper.

## Acknowledgments

We thank Juan C. de la Torre for sharing the LCMV rescue system with us. This research was supported in part by the Japan Agency for Medical Research and Development (AMED) (grant numbers, JP22fm0208101 and JP22ym0126815, to M.I.). We thank J. Ludovic Croxford, PhD, from Edanz (https://jp.edanz.com/ac) for editing a draft of this manuscript.

## Author contributions

M.H., A.T., and M.I. conducted experiments and acquired data; M.H. and M.I designed research studies and analyzed data; M.I. provided reagents and wrote the manuscript with inputs by all authors.

## Conflict of Interest

MH and MI are the inventors of a patent application related to the findings from this study. AY declares no competing interests.

