## Supplemental Figures for "An mRNA-LNP-based Lassa virus vaccine induces protective immunity in mice"

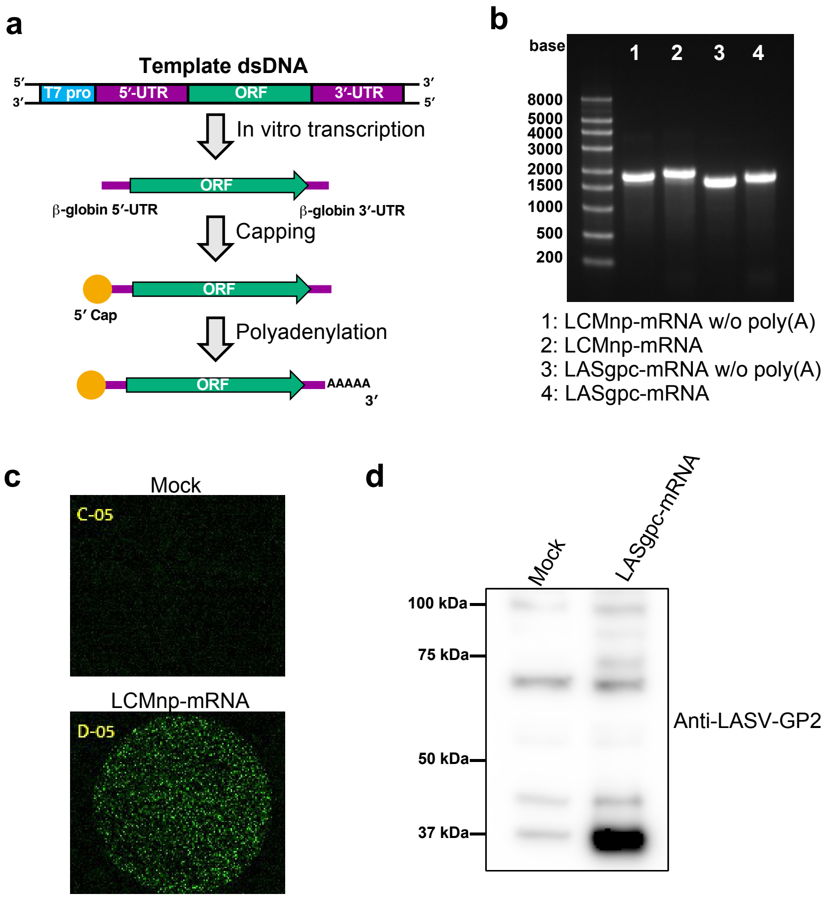


**Supplementary Fig. 1. Generation of in vitro transcribed (IVT) mRNAs expressing the LASV glycoprotein precursor (LASgpc-mRNA) and LCMV nucleoprotein (LCMnp-mRNA)**

**a** Schematic diagram of the method used to generate IVT mRNAs encoding viral protein open reading frames. The naked RNA containing the human β-globin 5'-UTR, a viral protein open reading frame, and the β-globin 3'-UTR was transcribed in vitro using a PCR-amplified DNA fragment as a template. A 5'-cap was added to the naked RNA and subsequently, the 3'-end was polyadenylated. **b** Agarose gel electrophoresis of IVT mRNAs with or without (w/o) poly(A). **c**, **d** HEK293T cells were transfected with 200 ng of LCMnp-mRNA (**c**) or LASgpc-mRNA (**d**), or mock transfected with Opti-MEM. At 24 h post-transfection, LCMnp levels in fixed cells were examined by an indirect immunofluorescence assay with a monoclonal antibody to LCMnp (**c**) and LASgpc levels in the clarified total cell lysate were examined by western blotting using a polyclonal antibody to LASV GP2 (**d**).

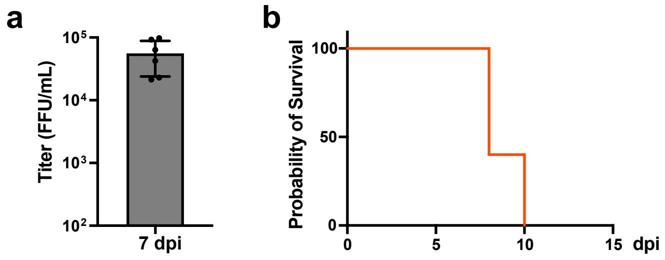


**Supplementary Fig. 2. Virulence of rLCMV/LASgpc$^{2m}$ in C57BL/6 mice**

**a** Eight-week-old C57BL/6 mice (n = 6) were inoculated i.v. with $10^6$ FFU of rLCMV/LASgpc$^{2m}$. At 7 dpi, viral titers in plasma were determined by an immunofocus forming assay. The presented data are the mean ± SD. **b** Eight-week-old C57BL/6 mice (n = 5) were inoculated i.c. with $10^3$ FFU of rLCMV/LASgpc$^{2m}$. Survival was monitored daily.

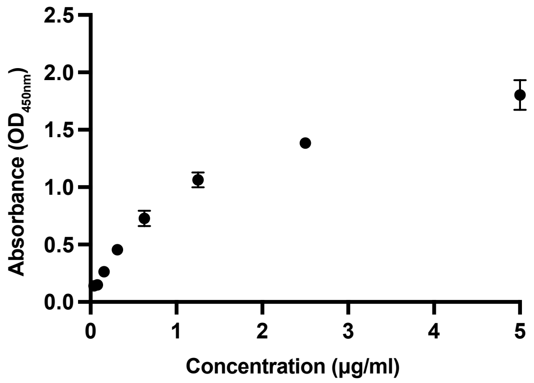


**Supplementary Fig. 3. Verification of an ELISA to detect LASgpc-specific antibodies**

The optical densities (ODs) of two-fold serial dilutions of anti-GP2 polyclonal antibody (PA5-117438) at 450 nm were determined by an ELISA developed in-house for the detection of anti-LASgpc-IgG.
